# Spatiotemporal functional interactivity among large-scale brain networks

**DOI:** 10.1101/2020.04.14.041830

**Authors:** Nan Xu, Peter C. Doerschuk, Shella D. Keilholz, R. Nathan Spreng

**Affiliations:** Wallace H. Coulter Department of Biomedical Engineering, Georgia Institute of Technology and Emory University, Atlanta, GA, United States; School of Electrical and Computer Engineering, Cornell University, Ithaca, NY, United States; Nancy E. and Peter C. Meinig School of Biomedical Engineering, Cornell University, Ithaca, NY, United States; Laboratory of Brain and Cognition, Montreal Neurological Institute, Department of Neurology & Neurosurgery, McGill University, Montreal, QC, Canada; Departments of Psychiatry and Psychology, McGill University, Montreal, QC, Canada; Douglas Hospital Research Centre, Montreal, QC, Canada; McConnell Brain Imaging Centre, Montreal Neurological Institute, McGill University, Montreal, QC, Canada

## Abstract

The macro-scale intrinsic functional network architecture of the human brain has been well characterized. Early studies revealed robust and enduring patterns of static connectivity, while more recent work has begun to explore the temporal dynamics of these large-scale brain networks. Little work to date has investigated directed connectivity within and between these networks, or the temporal patterns of afferent (input) and efferent (output) connections between network nodes. Leveraging a novel analytic approach, prediction correlation, we investigated the causal interactions within and between large-scale networks of the brain using resting-state fMRI. This technique allows us to characterize information transfer between brain regions in both the spatial (direction) and temporal (duration) scales. Using data from the Human Connectome Project (N=200) we applied prediction correlation techniques to four resting state fMRI runs (total TRs = 4800). Three central observations emerged. First, the strongest and longest duration connections were observed within the somatomotor, visual and dorsal attention networks. Second, the short duration connections were observed for high-degree nodes in the visual and default networks, as well as in hippocampus. Specifically, the connectivity profile of the highest-degree nodes was dominated by efferent connections to multiple cortical areas. Moderate high-degree nodes, particularly in hippocampal regions, showed an afferent connectivity profile. Finally, multimodal association nodes in lateral prefrontal brain regions demonstrated a short duration, bidirectional connectivity profile, consistent with this region’s role in integrative and modulatory processing. These results provide novel insights into the spatiotemporal dynamics of human brain function.

## 1. Introduction

Human brain function at rest (i.e. in the absence of task) is characterized by coherent and persistent patterns of regional interactions organized into spatially distributed, large-scale networks (Cole, Bassett et al. 2014). These intrinsic patterns emerge from repeated task-driven entrainment of afferent (input) and efferent (output) connections among brain regions (Stevens and Spreng 2014). Resting-state functional magnetic resonance imaging (fMRI), which examines the low frequency spontaneous fluctuations in blood oxygen level dependent (BOLD) signals (Biswal, Yetkin et al. 1995), is widely used to investigate the intrinsic functional architecture of the brain. Substantial progress has been made in delineating large-scale functional brain networks using resting-state functional connectivity (RSFC) (e.g., Biswal, Yetkin et al. 1995, Power, Cohen et al. 2011, Yeo, Krienen et al. 2011, Uddin, Yeo et al. 2019). The most commonly used method for measuring RSFC is to calculate a pairwise correlation coefficient between the low frequency BOLD signals of a set of priori determined brain regions of interest (ROIs). These correlation matrices may then be used to model the spatial topology of functional networks and sub-networks (Sporns 2011, Wig, Schlaggar et al. 2011).

In the last decade substantial efforts have been devoted to identifying the spatiotemporal structure of these RSFC-derived brain networks. However, comparatively less effort has been focused on identifying directional or ‘effective’ connectivity patterns. Several computational approaches have been suggested to explore directed intrinsic connectivity (e.g., Roebroeck, Formisano et al. 2005, Kim, Zhu et al. 2007, Blinowska, Trzaskowski et al. 2009, Chen, Glen et al. 2011, Xu, Spreng et al. 2017). In parallel, several studies have revealed that correlated, yet temporally asynchronous, patterns of BOLD signal may reflect the timing of information transfer in the brain (e.g., Mitra, Snyder et al. 2015, Yuste and Fairhall 2015, Goelman, Dan et al. 2017, Xu, Spreng et al. 2017). Together, these advances open the possibility of identifying not only the spatial properties of these intrinsic brain networks but also the direction and rate of information flow through this network architecture.

We have developed a novel statistical approach, prediction correlation (p-correlation, Xu, Spreng et al. 2017), to measure the direction and temporal profile of intrinsic functional connections. P-correlation provides a single analytical technique to characterize the strength, direction, and timing of connections simultaneously. We have validated the p-correlation approach using synthetic data for direction estimation and resting state fMRI (rs-fMRI) data to estimate connection strength (Xu, Spreng et al. 2017). As p-correlation decomposes asynchronous pairwise functional interactivity we are able to derive ‘direction’ as well as ‘duration’ information to characterize node to node connectivity profiles within and between networks. This approach is in contrast to symmetric estimates of RSFC as measured by standard Pearson correlation methods. To distinguish these measures from standard pairwise, symmetric *functional connectivity* (FC) measured by Pearson correlation, we refer to the functional interactivity measured by p-correlation as *directed functional connectivity* (directed FC), and *directed RSFC* for intrinsic network measures obtained at rest.

In the current study we apply p-correlation to rs-fMRI data to investigate the directed intrinsic functional connectivity of the human brain. To characterize the temporal flow of information through the networks, we first identified the duration (long versus short) of asynchronous connections across the brain and their spatial overlap with known intrinsic brain networks. Next we examined the asymmetry of the pairwise spatiotemporal functional interactivity. Our reasoning here was that p-correlation decomposes the pairwise functional interactivity to inward and outward information transfers, providing directional estimations. To our best knowledge, this is the first work in which the brain-wide spatial and temporal patterns of pairwise functional interactivities are jointly examined.

## 2. Methods

### 2.1. Participants and data preprocessing

The rs-fMRI data of 200 randomly-selected unrelated individuals was downloaded from the Human Connectome Project 500 Subjects + MEG2 dataset (https://www.humanconnectome.org/study/hcp-young-adult/document/500-subjects-data-release) (Marcus, Harwell et al. 2011, Van Essen, Ugurbil et al. 2012, Van Essen, Smith et al. 2013). This cohort of 200 participants was aged 22-36 years 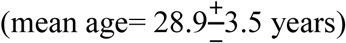, with 126 women.

The rs-fMRI data was collected at Washington University in St Louis and the University of Minnesota with the same sampling rate, *TR* = 0.72s. Each subject has 4 complete rs-fMRI scans, and each scan has 1200 time samples, i.e., *N*_*x*_ = 1200. Other acquisition parameters include TE=33.1ms, flip angle=52 degree, FOV=208×180 mm (RO × PE), Matrix 104×90 (RO x PE), Slice thickness=2.0 mm with 72 slices and 2.0 mm isotropic voxels, Multiband factor=8, Echo spacing=0.58ms, and BW=2290 Hz/Px.

The rs-fMRI data was first preprocessed following the procedures detailed in (Glasser, Sotiropoulos et al. 2013) to remove spatial distortions, motion artifacts, and reduce field bias. This procedure also registered the functional and anatomical images and normalized the data to standard space. The output fMRI data was then denoised by the ICA-FIX approach as described in detail in (Griffanti, Salimi-Khorshidi et al. 2014, Salimi-Khorshidi, Douaud et al. 2014). Finally, the preprocessed rs-fMRI BOLD timeseries were extracted from 333 ROIs (*N*_*ROI*_=333). These were centered in 333 discrete brain parcels provided in Gordon, Laumann et al. (2016). In the remainder of the paper, we refer the resulting dataset as the extracted rs-fMRI timeseries.

In addition to the extracted rs-fMRI timeseries, shuffled data and phase randomized data were also generated. The generation procedures and results are detailed in the Supplementary Methods and Results. The purpose of data shuffling or randomizing the phase of the data is to randomize the spatiotemporal dynamics of the original rs-fMRI timeseries. These results can then be used to contrast the spatiotemporal structure estimated from original data to validate the analytical approach under testing.

### 2.2. Mathematical description of the spatiotemporal functional interactivity

The spatiotemporal functional brain activity may be decomposed into two components, 1) the spatially directed FC and 2) the temporal durations of information transfer of this directed FC. In this section, we briefly review the mechanism of prediction correlation (p-correlation) introduced in Xu, Spreng et al. (2017) and describe how these two patterns are estimated from p-correlation.

Let *x*_*i*_ and *x*_*j*_ be the BOLD signals of *N*_*x*_ samples which come from the *i*th ROI and the *j*th ROI, respectively. P-correlation of the ordered pair (*x*_*i*_, *x*_*j*_) involves several computational steps. The first step estimates the duration of information transfer from the *i* th ROI and the *j* th ROI. Specifically, the output signal *x*_*j*_ is predicted from the input signal *x*_*i*_ through a linear time-invariant causal dynamic model that characterizes the information transfer, and the prediction, denoted by *x*_*j*|*i*_, has the form

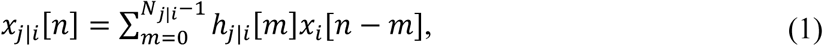

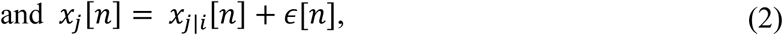

here *h*_*j*|*i*_ is a *N*_*j*|*i*_-length impulse response vector which has the first potentially non-zero term occurred at time of 0s, and *ϵ* is the prediction error. The current sample of the output signal *x*_*j*_ is always effected by *N*_*j*|*i*_ time lags of the input signal *x*_*i*_. Hence, the duration of the directed information transfer has *TR* · *N*_*j*|*i*_ seconds. The optimal solution for *N*_*j*|*i*_ is determined by Bayesian information criterion (BIC) through testing from 1 up to 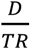 samples to minimize the prediction error. In this study, we restrict the temporal window for directional influence between ROIs to be no more than 15s, i.e., *D* =15s.

The second step estimates the strength of the RSFC between the ordered pair (*x*_*i*_, *x*_*j*_), denoted by p-corr(*x*_*i*_, *x*_*j*_). Specifically, a correlation between the predicted signal *x*_*j*|*i*_ and the original BOLD signal *x*_*j*_ is computed.

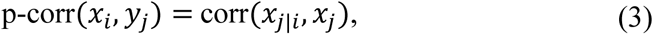

Note that if the signal *x*_*j*_ and its prediction *x*_*j*|*i*_ are significantly correlated, i.e., the result of Eqn. (3) is high, then the likelihood of a directed information transfer as modeled in Eqn. (1-2) is high. The significance of the p-correlation can be determined by a p-value test. The p-value at which the null hypothesis of zero p-correlation is rejected (probability whose small value indicates a significant p-correlation) can be computed by following the procedure in Press (2007).

As described in Xu, Spreng et al. (2017), p-correlation is able to replicate previously observed modular network structures in resting brain (Power, Cohen et al. 2011). Thus, p-corr(*x*_*j*_, *x*_*i*_) can be used to evaluate the strength of RSFC from *i*th ROI to the *j*th ROI. However, unlike the standard correlation, p-correlation is asymmetric between the two signals *x*_*i*_ and *x*_*j*_. Specifically, p-corr *x*_*i*_, *x*_*j*_ ≠ p-corr (*x*_*j*_, *x*_*i*_) and *N*_*j*|*i*_ ≠ *N*_*i*|*j*_. The asymmetry between p-corr (*x*_*i*_, *x*_*j*_) and p-corr(*x*_*j*_, *x*_*i*_) leads to a directed graph between two brain regions in the spatial scale. In addition, the separate estimations for *N*_*i*|*j*_ and *N*_*j*|*i*_ characterizes the interactions in the temporal scale.

For the whole-brain fMRI timeseries, two *N*_*ROI*_ × *N*_*ROI*_ asymmetric matrices were generated by p-correlations. One is a connectivity matrix including p-correlation values (e.g., p-corr(*x*_*i*_, *x*_*j*_) from Eqn. (3)). The other is a duration matrix including the duration of the information transfer (e.g., *TR* · *N*_*j*|*i*_) estimated from Eqn. (1-2).

### 2.3. Analytical procedures

In this section, the analytical procedures for investigating the spatiotemporal structure estimated from p-correlation are described. First, the spatial pattern and the temporal pattern of the functional interactivity were determined by p-correlation using the extracted rs-fMRI timeseries. Specifically, a directed FC map and a duration map was determined by averaging the connectivity matrices and the duration matrices across all scans of all 200 subjects, respectively. To validate the spatiotemporal patterns estimated by p-correlation on the extracted rs-fMRI timeseires, the directed FC map and duration map were contrasted with the results of the shuffled data and of the phase-randomized data (see Supplementary Methods and Results for more details). Seven functional networks provided in Yeo, Krienen et al. (2011) were used in the remainder analysis, which include Somatomotor (SM), Visual, Dorsal, Ventral, Default, Frontal parietal (FP), and Limbic networks (see Supplementary Methods and Results for more details).

Next, we investigated the directed RSFC with the longest and shortest duration of information transfer in the human brain. In particular, the longest durations of information transfers determined by the top 4% values in the duration map were compared with the strongest connections as determined by the top 4% values in the functional map. The directed RSFC with the shortest durations of information transfer were determined by the smallest 4% nonzero values in the duration map which has p-value<0.05 in the directed FC map. The functional interactivity among networks was evaluated for the directed FC map as well as the duration maps, and is described in the Supplementary Methods and Results.

Finally, we assessed the asymmetry between the inward and outward information transfer for each ROI as well as for every functional network. In particular, the upper off-diagonal entries were subtracted from their lower off-diagonal counterparts (referred as the afferent-efferent, or inward-outward differences in the remainder of the paper) in both the directed FC map and duration map. The mean and standard deviation of the inward-outward differences were computed. In addition, due to the novelty of the duration map estimation, its inward and outward asymmetries were examined in more detail. Specifically, for each of the 333 ROIs, the row mean and the column mean of the duration map were computed to give the average durations of the inward and outward information transfers, respectively. Furthermore, the duration map was partitioned into 7 × 7 subblocks, in which the diagonal subblocks includes the connectivity within functional networks and the off-diagonal subblocks includes the connectivity between different pairs of networks. The average duration of the inward (outward) information transfer for the *i*th functional brain network was determined by computing the mean and standard deviation of entries in the *i*th column (row) partition excluding the *i*th diagonal subblock of the duration map, for *i* =1, …, 7.

## 3. Results

### 3.1. Long duration rs-fMRI network connectivity

The directed FC map and duration maps estimated from p-correlation is shown in Fig. 1 (a) and Fig. 1 (b), respectively. Note that global signal regression was not applied to rs-fMRI data in the current study due to the current controversial effects (Murphy and Fox 2017). Hence, p-correlation values are mostly positive in Fig. 1 (a). As shown, organized patterns are demonstrated in both the directed FC and duration maps. In particular, the strong connections are concentrated in the diagonal blocks, suggesting that connections within each functional network on average are stronger than the ones between this network and the other networks. Similar to the directed FC map, the high values of the duration map are also concentrated within the diagonal blocks, with high overlap between strong and long duration functional connections. The correlation between the directed FC map and the duration map was computed as 0.72. The top 4% of values in either of these two maps are further shown in Fig. 1 (c) and (d), respectively. The strongest directed connections and the longest durations occur primarily within-networks, including the SM, visual and ventral systems. Longer durations were also observed between networks (e.g. SM - Visual networks). On the other hand, directed functional interactivities within the limbic system have much lower values in both maps comparing to other functional networks.

**Figure 1.**
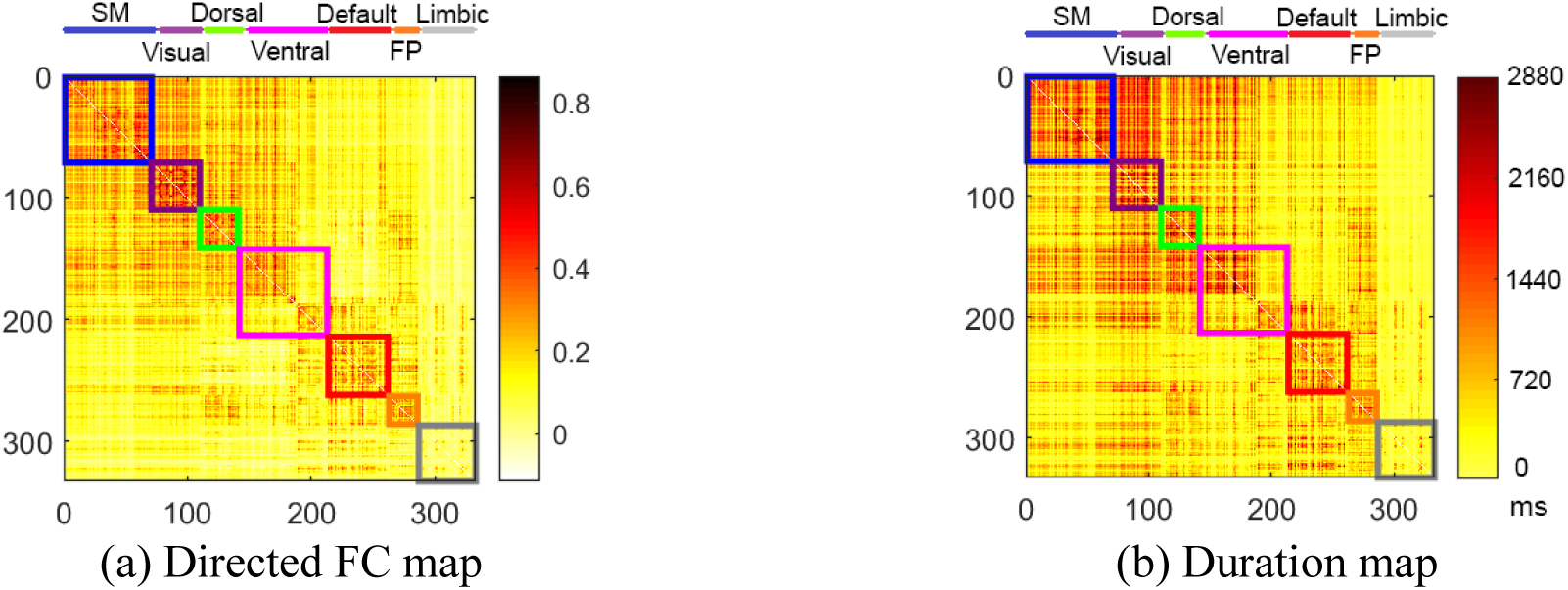
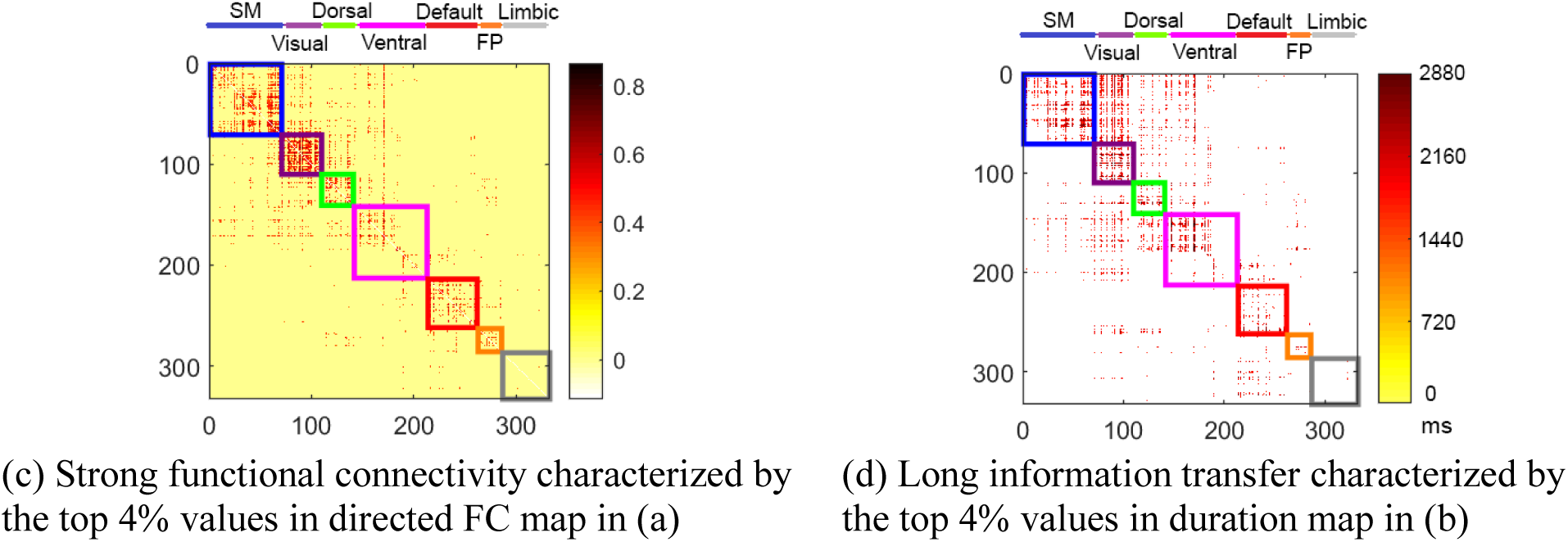
The directed functional connectivity map and the duration map of resting-state functional networks determined by p-correlations. In these matrices, ROIs along the rows propagate information to the ROIs along the columns. The strongest directed FC and the longest information transfers are shown in (c) and (d), respectively. Seven colored diagonal blocks in each matrix depict the seven different networks including SM (blue), Visual (purple), Dorsal (green), Ventral (pink), Default (red), FP (orange), Limbic (grey).

### 3.2. Short-duration rs-fMRI network connectivity

The duration map and the corresponding directed RSFC map with short information transfers as described in Section 2.3 are shown in Fig. 2(a) and (b), respectively, and displayed in Fig. 2 (c). The smallest 4% nonzero values in the duration map (without considering the p-value for p-correlations) and the corresponding connectivity map are shown in Figure S6 of Supplementary Methods and Results.

**Figure 2.**
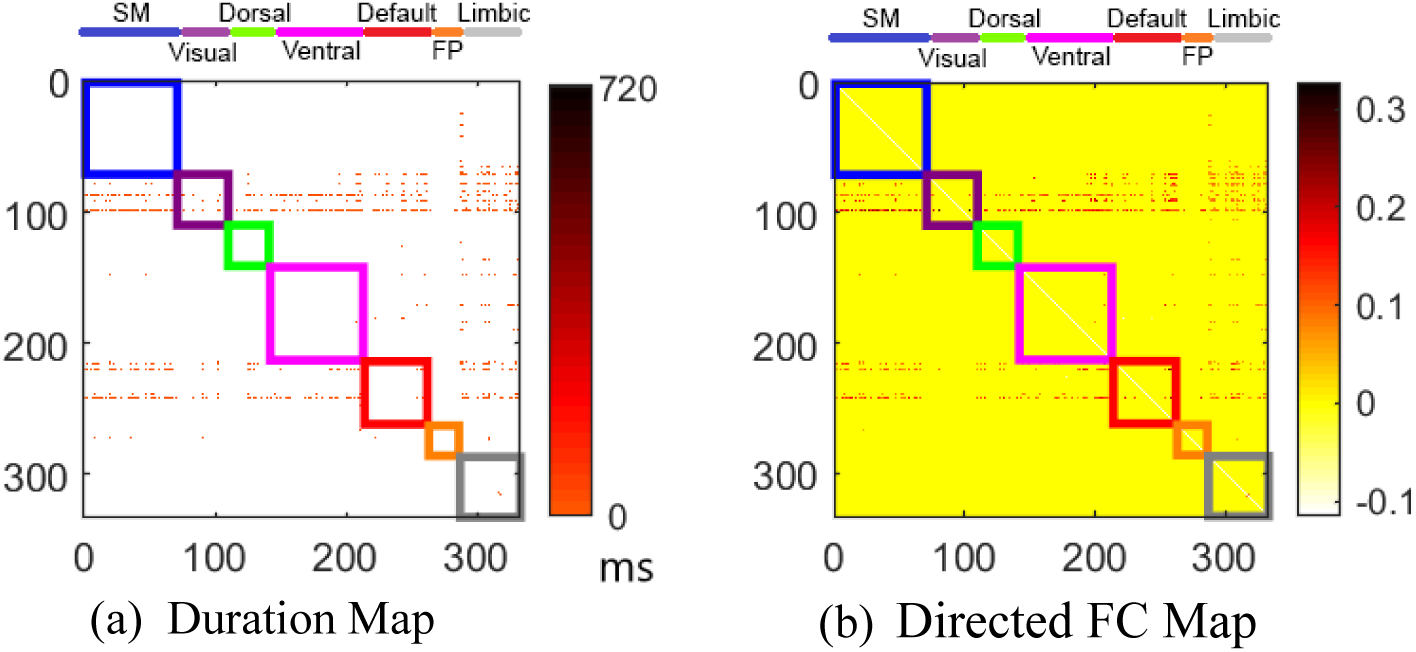
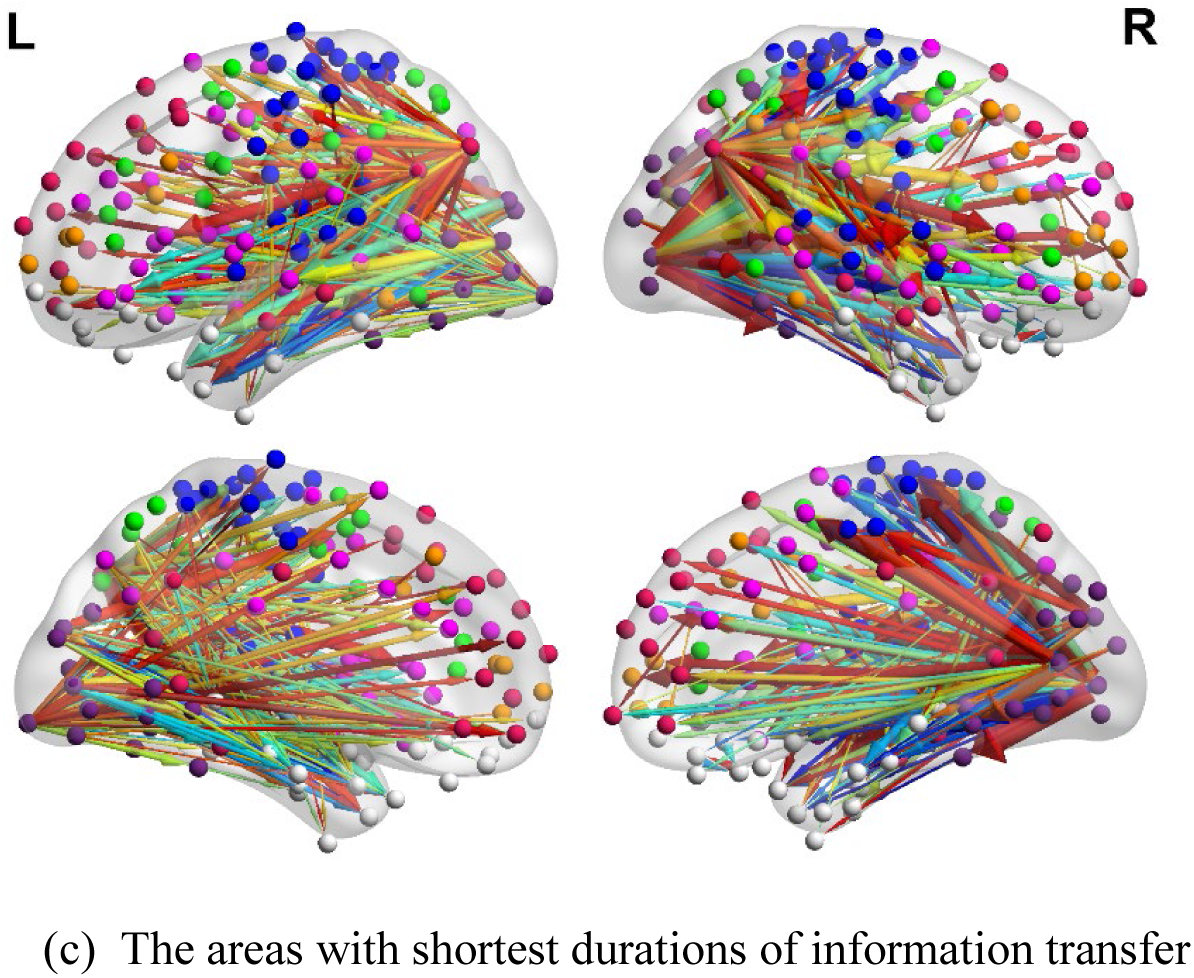
Short information transfer, characterized by the shortest 4% nonzero *N*_*j*|*i*_ values and p-correlations with p-value<0.05. The duration map (a) and the directed functional connectivity map (b) of the short information transfer in rs-fMRI. In both matrices, ROIs along the rows propagate information to the ROIs along the columns. Seven colored diagonal blocks in (a) and (b) depict the seven different networks as described in the legend (right). (c) The areas with the shortest information transfer are displayed. The color of the arrow from cool to warm colors represents the duration from short to long. The thickness of the arrow represents the strength of the connectivity. In total, 1099 directed functional connections are demonstrated. Seven colored diagonal blocks in (a) and (b) as well as the node color in (c) depict the seven different networks including SM (blue), Visual (purple), Dorsal (green), Ventral (pink), Default (red), FP (orange), Limbic (grey).

Node degree is determined by the number of network connections (Power, Schlaggar et al. 2013). High degree ROIs are those with extensive functional connections relative to other brain networks. These may serve as information processing hubs (Power, Schlaggar et al. 2013).

As observed in Fig. 2, the short duration connections are dominated by high degree ROIs. Further, rows with dense dotted lines of the two matrices shown in Fig. 2 suggest that these high degree, short duration hubs primarily are dominated by efferent connections to other ROIs and networks, and we have labeled these as “outward brain hubs.” However, the columns with less dense dotted lines in the two matrices in Fig. 2 also identify short duration, high degree hubs that show a predominantly afferent, or inward, connectivity pattern, and these hubs are referred to as “inward brain hubs.”

High-degree outward and inward brain hubs are displayed on cortical maps in Figure 3 (a) and (b), respectively. Efferent or outward brain hubs with the highest degree are located in the visual system and the default system, in particular, the angular gyrus. Specifically, the top 5 outward brain hubs (Figure 3 (a)) are #263 in Visual (degree 181), #259 in Default (degree 121), #140 in Visual (degree 110), #94 in Default (degree 88), #6 in Default (degree 63)^1^. Afferent (or inward) brain hubs with high degrees are primarily located in the limbic system. The top 14 inward brain hubs (Figure 3 (b)) include #178 in Limbic (degree 31), #19 in Limbic (degree 27), #135 in Limbic (degree 21), #18 in Limbic (degree 18), #11 in Limbic (degree 18), #314 in Limbic (degree 17), #144 in Limbic (degree 17), #300 in Limbic (degree 15), #118 in Limbic (degree 15), #312 in Limbic (degree 14), #142 in Limbic (degree 14), #304 in Limbic (degree 13), #296 in Limbic (degree 13), and #295 in Default (degree 13)^1^. Both afferent and efferent hubs show high hemispheric symmetry. The efferent hubs have higher degree values than the hubs that demonstrate a more afferent, or inward, pattern of directed connectivity.

**Figure 3.**
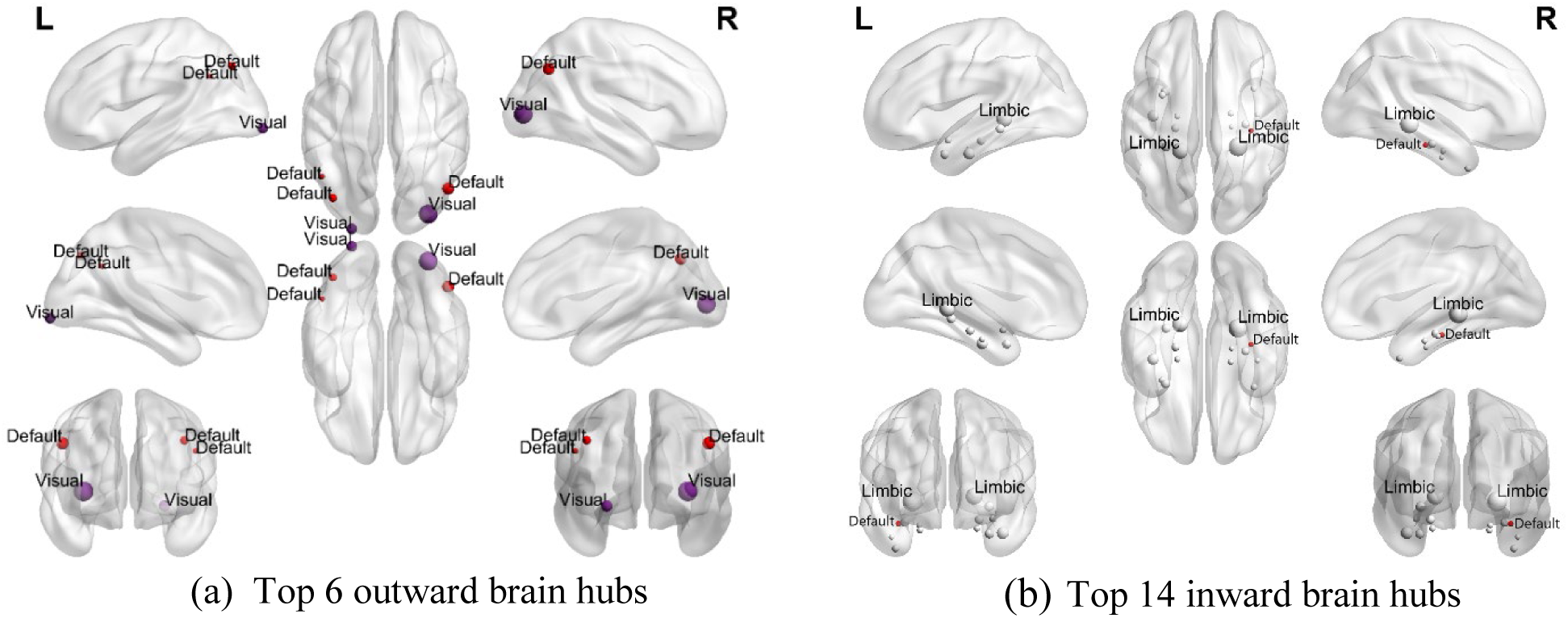
Brain hubs displaying short information transfer in function networks. The node color represents the brain system of visual (purple), default (red), and limbic (white). The node size is proportional to the hub degree in each subfigure.

### 3.3. Asymmetries between the inward and outward information transfer

Asymmetries between inward and outward information transfer exist in both the directed FC map and duration map for every pair of ROIs (Fig. 1) as well as for every pair of functional networks (Fig. S2). To visualize the asymmetries in detail, the information of the connectivity (Fig. 1 (a)) and duration (Fig. 1 (b) maps are shown in Fig. 4. This figure displays the top 100 strongest directed connections within, going out of, and coming into each of the seven functional brain networks. The top 1% strongest directed connections within, going out of, and coming into each of the seven functional brain networks are in Fig. S6 in the Supplementary Methods and Results. These figures clearly show that the inward-outward asymmetries are much more pronounced for the duration versus the direction estimates, and this pattern is observed for every network and even for every ROI. Statistically, each pair of ROIs has an average of inward-outward differences of 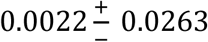 in directed RSFC strengths, and of 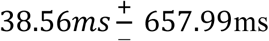 in durations. Each pair of functional networks has an average of outward-inward differences of 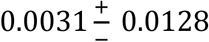 in directed RSFC strengths, and of 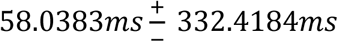 in durations.

**Figure 4.**
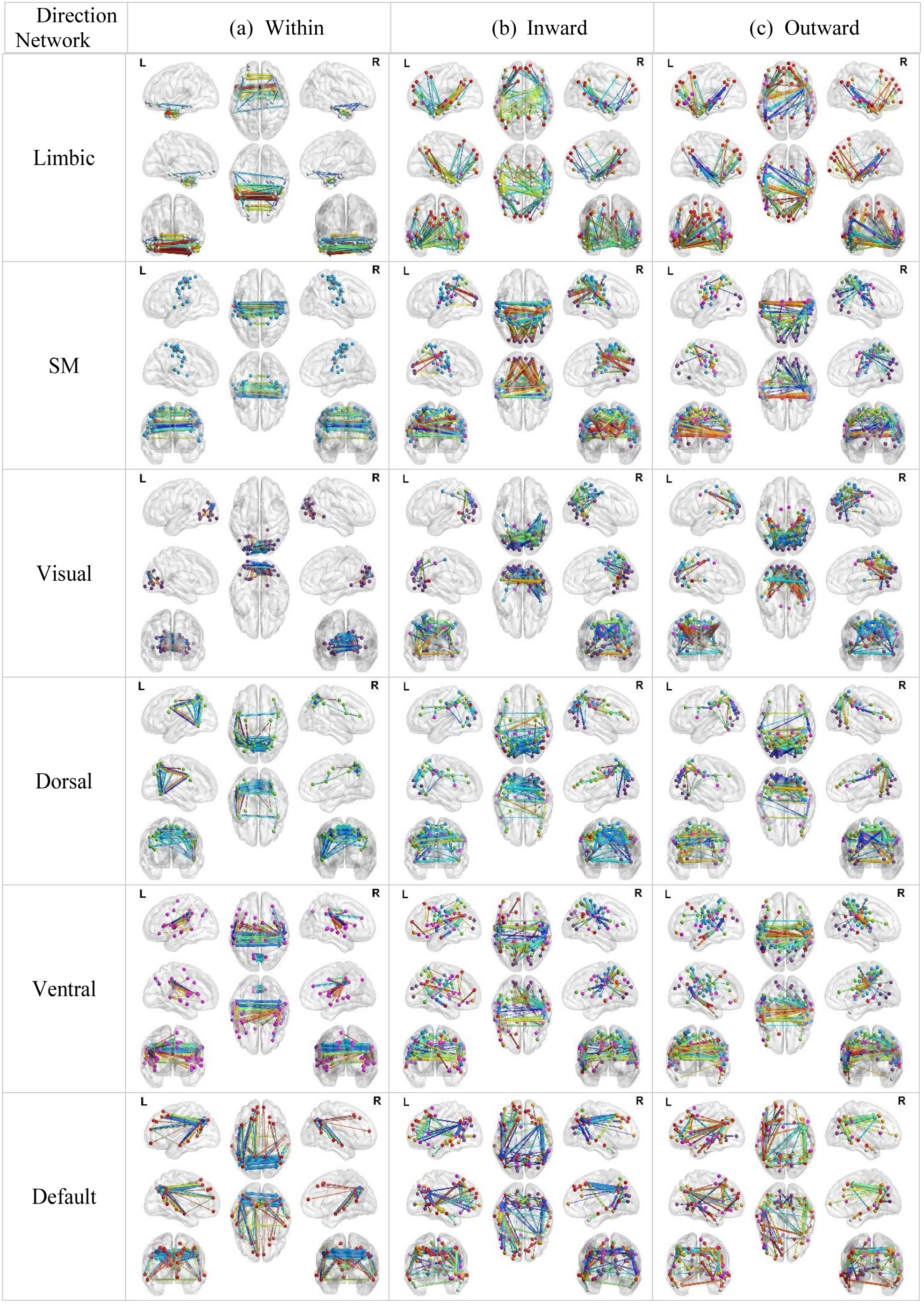
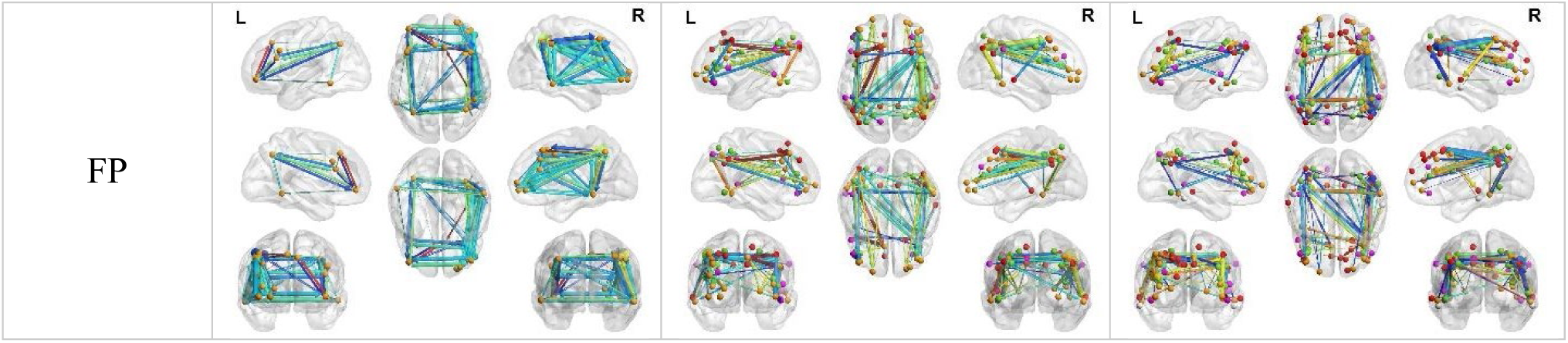
Spatiotemporal functional interactivity in resting state brain networks. The top 100 strongest connections within (a), coming into (b), and going out of (c) each of the seven functional brain networks are visualized by a modified version of BrainNet Viewer toolbox (Xia, Wang et al. 2013). The duration and the strength of the information are jointly displayed. In particular, the color of the arrow from cold to hot represents the duration from short to long. The thickness of the arrow represents the strength of the connectivity. So, stronger and longer connectivity has a thicker and warmer colored arrow. ROIs in each of the seven networks share the same network color as shown in the legend of Fig. 3 (right).

We next evaluated the asymmetries in the duration maps in more detail. Average durations for inward and outward information transfer for each network as well as their differences are presented in Table 1. Visual and Limbic networks show the greatest inward-outward differences in duration. In contrast, the Ventral and Default networks have more similar inward and outward connections in duration. The duration of information transfers for the SM, Ventral and Limbic networks are generally shorter for inward versus outward connections, whereas all other networks show the reverse pattern, with shorter duration outward versus inward connections. Overall, the Limbic, Ventral and Default networks have the shortest inward information transfers, whereas FP, Default and Visual networks have the shortest outward information transfers.

**Table 1.** The average duration (ms) of the inward and outward information transfer and their differences for each network.

| Network | Direction | Inward | Outward | Inward-Outward Differences |
| --- | --- | --- | --- | --- |
| SM |  | 1205<sup>+</sup><sub>345</sub> | 1371<sup>+</sup><sub>517</sub> | -166<sup>+</sup><sub>643</sub> |
| Visual |  | 1581<sup>+</sup><sub>527</sub> | 1130<sup>+</sup><sub>370</sub> | 451<sup>+</sup><sub>659</sub> |
| Dorsal |  | 1355<sup>+</sup><sub>442</sub> | 1186<sup>+</sup><sub>392</sub> | 169<sup>+</sup><sub>565</sub> |
| Ventral |  | 1198<sup>+</sup><sub>396</sub> | 1313<sup>+</sup><sub>457</sub> | -115<sup>+</sup><sub>586</sub> |
| Default |  | 1199<sup>+</sup><sub>389</sub> | 1071<sup>+</sup><sub>301</sub> | 127<sup>+</sup><sub>502</sub> |
| FP |  | 1257<sup>+</sup><sub>296</sub> | 1063<sup>+</sup><sub>295</sub> | 195<sup>+</sup><sub>402</sub> |
| Limbic |  | 862<sup>+</sup><sub>150</sub> | 1218<sup>+</sup><sub>355</sub> | -355<sup>+</sup><sub>389</sub> |

Average durations of the inward and outward information transfers for each ROI are displayed in Fig. 5. Examining specific asymmetry patterns, the occipital lobe (Visual and Dorsal ROIs) displays the greatest temporal asymmetry with the longest inward and shortest outward durations. In contrast, the Limbic ROI shows the reverse pattern, with the shortest duration inward connection but the second longest outward connection. The Ventral and SM ROIs show a somewhat more balanced duration pattern, albeit with overall longer outward than inward durations. ROIs within the PFC show the most equivalent inward and outward pattern, with nearly identical durations (∼1300 ms). Finally, as observed in Fig 5, there are nodal differences in duration asymmetry within networks. For example, the Default network precuneus ROI shows a longer inward and shorter outward duration pattern, whereas the medial prefrontal ROIs of the default network shows shower inward and shorter outward duration patterns.

**Figure 5.**
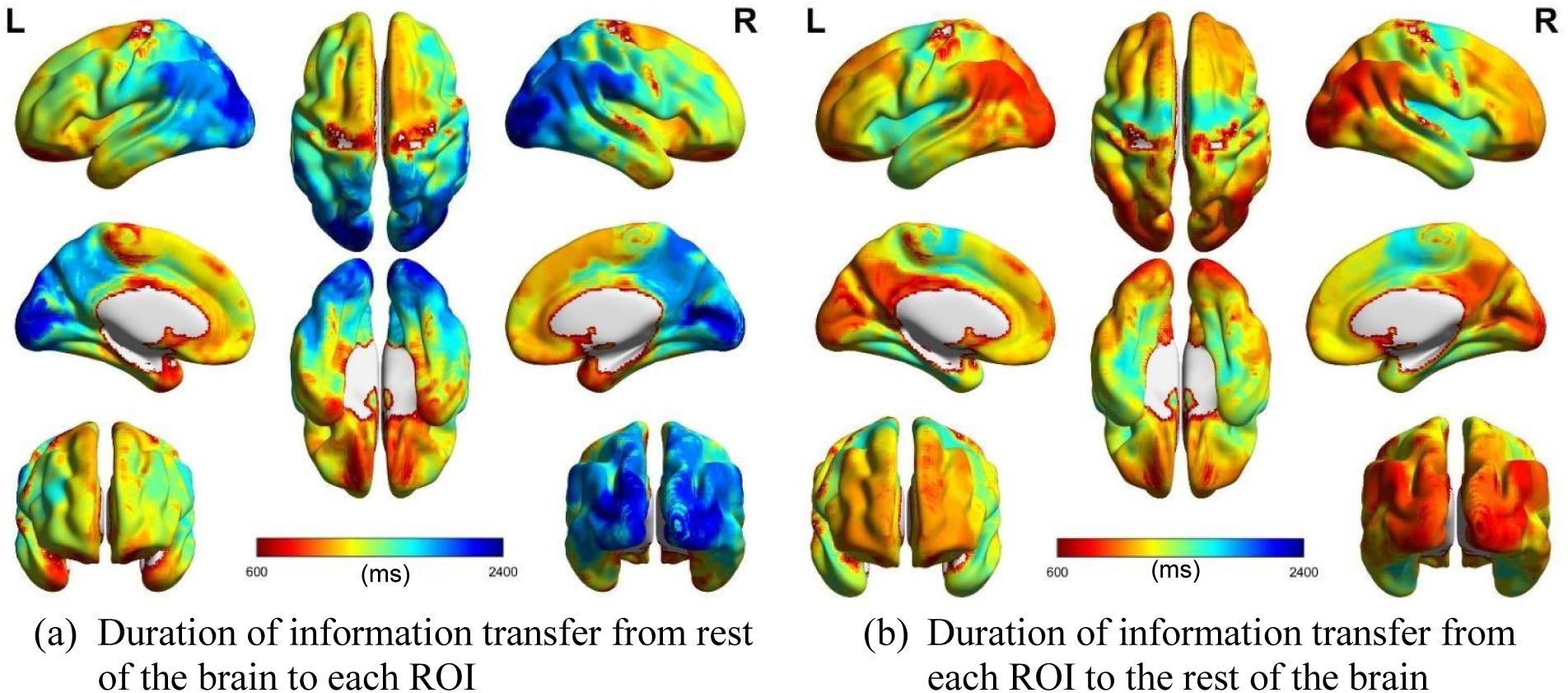
The average duration (ms) of the information transfer coming from and going into each of the 333 ROIs provided in Gordon, Laumann et al. (2016). The shortest information transfers have warm colors, while the longest information transfers have cool colors. Both inward and outward information transfers are displayed in the same colormap.

## 4. Discussion

Here we used a novel approach, p-correlation, to estimate the direction and duration of information transfer within a resting-state fMRI dataset. The p-correlation method decomposes the traditional pairwise functional interactivity into inward and outward functional connectivity. The connection strength as well as the duration of the information transfer are estimated for each direction. This method enables us to measure the direction (inward versus outward) and the duration (long versus short) connectivity profiles for all ROI-ROI pairs. The resulting whole-brain, directed functional connectivity and duration maps revealed distinct network and regional patterns of spatiotemporal connectivity within the resting brain.

### Long vs. short duration connections

Long duration information transfers are observed within the somatomotor, visual and dorsal attention networks. These long duration connections are highly overlapping with the stronger directed connections (Fig. 1). The shortest information transfers, on the other hand, are driven by multiple high-degree outward brain hubs in the visual and default systems (i.e., rows with dense dotted lines in Fig. 2) as well as several moderate high-degree inward brain hubs in the hippocampus (i.e., columns with less dense dotted lines in Fig. 2). The outward hubs, that segregate information from visual or default system to the other cortical regions (as shown in Fig. 2 (c) and Fig. 3 (a)), in particular are consistent with the previous findings of functional hubs in cuneus of the visual system (Tomasi and Volkow 2011) as well as the angular gyrus of default system (Andrews-Hanna, Smallwood et al. 2014). Our results further suggest that the role of these high-degree brain hubs is actively propagating the information to, instead of receiving the information from, other brain regions in a short temporal scale. In addition, the inward brain hubs in the limbic system which aggregate the information from the cortex (as shown in Fig. 2 (c) and Fig. 3 (b)), is consistent with the idea that resting state brain function supports the memory consolidation (Miall and Robertson 2006, Stevens, Buckner et al. 2010, Tambini, Ketz et al. 2010, Gordon, Breeden et al. 2014, Stevens and Spreng 2014).

### Inward and outward asymmetry of the information transfer

In contrast to many other lag-based analytical approaches which assume the temporal lags are pairwise identical, e.g., (Mitra, Snyder et al. 2015, Raut, Mitra et al. 2020), we assume the functional interactivity can be pairwise asymmetric not only in the connection strength but also in the duration of the information transfer. In other words, ongoing information transfer in both forward and backward directions may simultaneously present in both spatial and temporal scales. Our findings, however, revealed only modest differences in the strength of directed pairwise connections (i.e. inward versus outward). Yet, there were striking asymmetries observed in the duration of these directed connections. This observation, as shown in Fig. 4, is persistent for every network and even for most ROI-ROI pairs. Furthermore, Fig. 4, which demonstrates the strongest directed connections within, going out of, and coming into each of the seven functional brain network, provides a detailed description of the spatiotemporal functional interactivities on brain cortex which may further our understanding of human cognition. In the end, the brain-wide duration of the information transfer was examined in more detail. As shown in Fig. 5, visual processing regions, including precuneus, show longer duration of inward, or afferent, connections relative to shorter duration of outward, or efferent, connections. In contrast, regions of the limbic system show longer duration of outward, or efferent, connections relative to shorter duration of inward, or afferent connections. This is consistent with its role in consolidation processes, particularly involving medial temporal lobe regions (p. 313, Purves, Cabeza et al. 2008, Schwab, Harbord et al. 2018). The prefrontal cortex on the other hand displayed inward and outward connections that were similar in duration, consistent with the role of this region as a heteromodal hub with dense, reciprocal connection to the rest of cortex (Mesulam 1998, Baars and Gage 2013).

### Directed information transfer within and between networks

Static correlation methods have robustly revealed the macroscale intrinsic network architecture of the human brain (e.g., Uddin, Yeo et al. 2019). These networks of RSFC are built from symmetrical correlation matrices. P-correlation extends this approach, allowing for an asymmetric matrix of connectivity. This approach provides novel insights into within network connectivity, how connections enter a network (inward connections), and how a network communicates with the rest of the brain (outward connections). Internetwork nodes of the default, frontoparietal control and dorsal attention networks differentially connect networks to each other (Spreng, Sepulcre et al. 2013, Dixon, De La Vega et al. 2018). The present work determines, for the first time, patterns of whole brain connections, and temporal dynamics, between networks along with the outgoing and incoming direction of information flow (Figure 4). The relative differences in the duration of incoming and outgoing connections may be of particular importance to understanding each network’s cognitive contributions to information processing along a hierarchical gradients of information processing (Margulies, Ghosh et al. 2016). RSFC network architectures closely align with task-based connectivity (Smith, Fox et al. 2009). Thus the patterns of information flow within and between networks reported in the resting state may reflect task-driven connectivity patterns implicated in cognitive performance. Short duration, synchronous information flows may serve as a putative mechanism underlying flexibility in network configurations and associated cognitive operations. Investigating associations between asynchronous and synchronous connectivity patterns and specific cognitive abilities will be an important direction for future research.

### Strengths and limitations

Comparing to the existing regression model-based methods for estimating the effective connectivity, p-correlation has several unique strengths. For example, in contrast to the widely used Granger causality statistic which depends on the assumptions of the errors of the autoregressive model to be Gaussian, p-correlation is based on just the sample variance of the prediction error and does not require a Gaussian assumption on the noise. This is advantageous if the BOLD signals lack Gaussian structure. Indeed, p-correlation has performed well on both synthetic and experimental fMRI data (Xu, Spreng et al. 2017). In addition, a significant merit of p-correlation, comparing to Granger causality or other wildly used multivariate vector autoregressive modeling (MVAR) (Kus, Kaminski et al. 2004, Wilke, Ding et al. 2008, Ligeza, Wyczesany et al. 2016) which measures the strength of *effective connectivity* purely by the weights of impulse response, is that p-correlation determines the strength of *functional interactivity* (Equation 3) that can be used to detect modular organizations of the brain (Xu, Spreng et al. 2017). Despite these advantages, the duration of information transfer estimated by p-correlation is particularly subject to sampling variability. As such, only spontaneous directed brain activities in resting brain that are longer than 1 *TTTT* can be detected by the method. Nevertheless, p-correlation is a measure which is adaptable to any type of timeseries, which enables us to explore the neural information processing in detail with electrophysiological data such as EEG and MEG that have finer timescales and higher SNR.

### Conclusions

As reviewed above, the p-correlation method provides novel insights into the spatiotemporal connectivity patterns in the resting brain. To our knowledge this is the first method to concurrently estimate the strength, direction and duration of resting-state state connectivity. As such, this method opens new avenues for understanding not only large-scale coherence patterns in the resting brain but also their temporal dynamics. This latter aspect, revealed by p-correlation, may serve as a proxy for investigating the nature of information flow through the intrinsic functional architecture of the brain. These insights in turn open a novel frontier for investigating associations between the intrinsic functional connectivity of the brain and human behavior.

## Supporting information

Supplemental Methods and Results

## Acknowledgement

“Data were provided [in part] by the Human Connectome Project, WU-Minn Consortium (Principal Investigators: David Van Essen and Kamil Ugurbil; 1U54MH091657) funded by the 16 NIH Institutes and Centers that support the NIH Blueprint for Neuroscience Research; and by the McDonnell Center for Systems Neuroscience at Washington University.” NX and SDK thank National Science Foundation (grant 1533260) for funding support. PCD thanks National Science Foundation (grant 1217867) for funding support. RNS thanks the support by the Natural Sciences and Engineering Research Council of Canada and Canadian Institutes of Health Research, and the Research Scholar support by Fonds de recherche du Québec – Santé.

## Footnotes

1 The # is consistent with the order of the 333 ROIs provided in Gordon, Laumann et al. (2016).

## Notes

### Competing Interest Statement

The authors have declared no competing interest.

