## Supplemental Methods and Results for "Spatiotemporal functional interactivity among large-scale brain networks"

### 1. Validation of p-correlation on shuffled data and phase-randomized data

#### 1.1. Data Generation

Data shuffling aims to destroy the spatiotemporal dynamics of the extracted rs-fMRI timeseries. Specifically, the extracted rs-fMRI timeseries of the $i$thROI was selected from a random subject at a random scan for $i = 1, \dots, N_{ROI}$. So, a set of shuffled data with 333 ROIs was generated for a whole brain.

The phase-randomized data were generated through shuffling the phase of the extracted rs-fMRI timeseries in the Fourier domain while preserving the power spectral magnitude and the correlational nature of the timeseries (Prichard and Theiler 1994, Majeed, Magnuson et al. 2011). This procedure would hence disturb the original temporal dynamics of the extracted rs-fMRI timeseries. In this study, a subset of 5 subjects out of the whole cohort are selected and following the procedure of (Majeed, Magnuson et al. 2011), phase randomized timeseries were generated for each ROI of every subject at each scan.

#### 1.2. Results

To validate the spatiotemporal structure estimated by p-correlation, the functional connectivity map and duration map estimated from the extracted rs-fMRI timeseries were compared with the ones computed from the shuffled data (Fig. S1) as well as the phase-randomized data (Fig. S2). In contrast to the results in Fig. 1, no organized patterns nor symmetric patterns are detected in either functional connectivity or duration information in Fig. S1. Numerically, the functional connectivity map has a mean of $0.0005 \pm 0.0414$, and the duration map has a mean of

$8.94 \pm 139.82$ ms. In the duration map, specifically, 98.96% of the pairwise connectivity has the information transfer with a duration of 0ms.

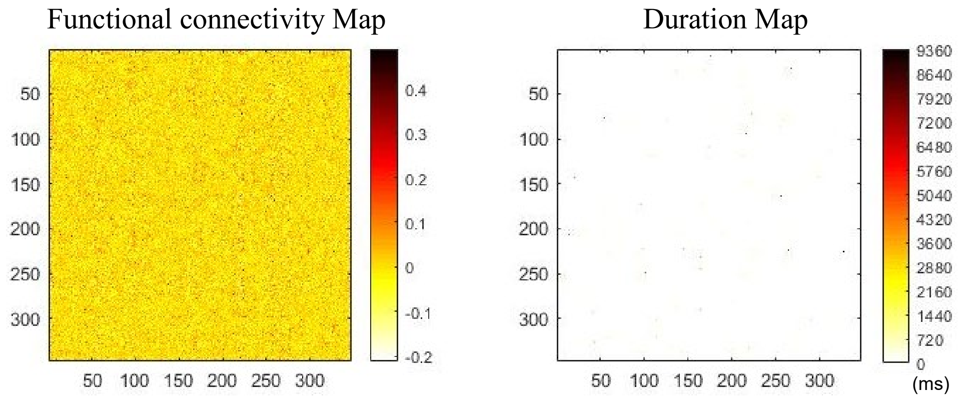


Figure S1 The connectivity map and the duration map of resting-state functional networks determined by p-correlations on shuffled data.

The phase-randomized data results is contrasted to the results of the extracted rs-fMRI timeseries on the same five subjects. As shown in Figure S2, block organized patterns on both the connectivity and duration maps are detected by p-correlation on the extracted rs-fMRI timeseries, and the block organized patterns are reproducible across different subjects. In contrast, no such reproducible block patterns are demonstrated in the results of the phase randomized data in both the functional connectivity and the duration. Numerically, the average connectivity map and the average duration map of the extracted rs-fMRI timeseries across all scans and subjects has a mean of $0.1561 \pm 0.1193$, and a mean of $1161.30 \pm 632.63$ ms, respectively. Comparatively, the average result of the phase randomized data has a mean of $0.0008 \pm 0.0219$ for the functional connectivity and a mean of $729.88 \pm 96.09$ ms for durations. In the duration map, specifically, 88.79% of the pairwise connectivity has the information transfer with a duration of 0ms.

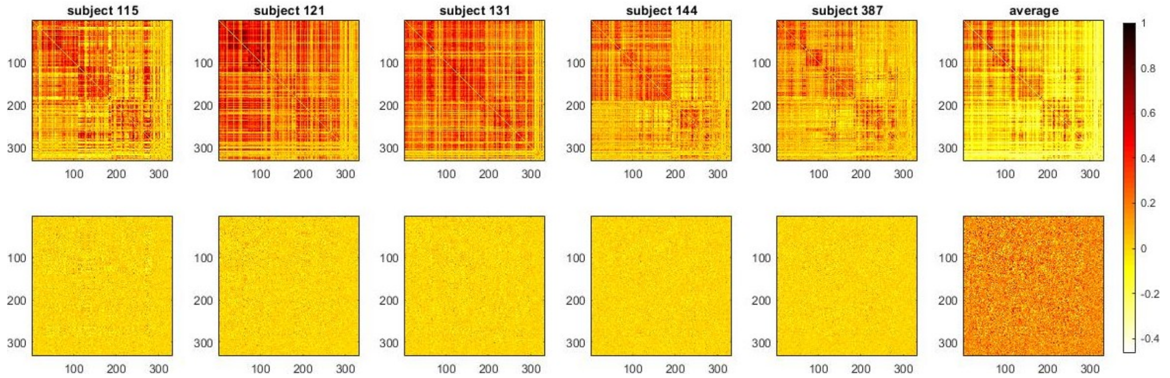


(a) Functional connectivity map determined by p-correlation on the extracted rs-fMRI timeseries (top panel) and the random shuffled data (bottom panel).

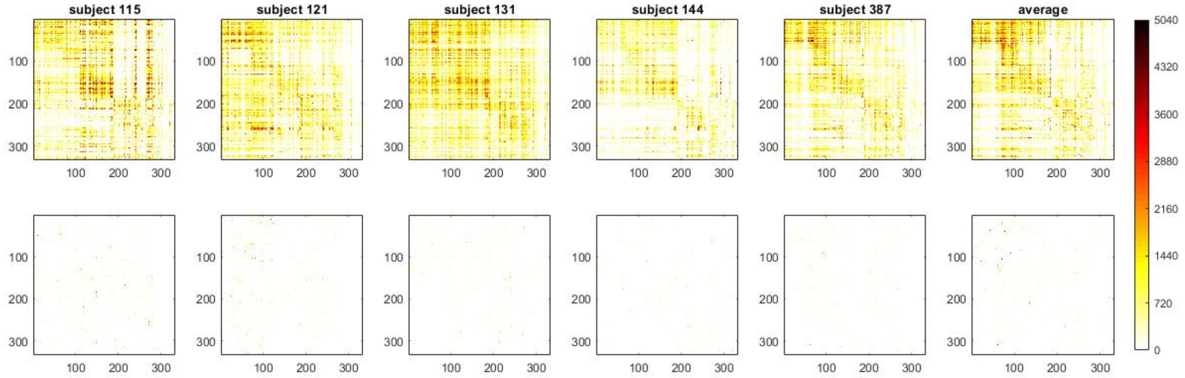


(b) Duration map determined by p-correlation on the extracted rs-fMRI timeseries (top panel) and the random shuffled data (bottom panel).

Figure S2 The connectivity map and the duration map of resting-state functional networks estimated by p-correlations on phase randomized data for five randomly selected subjects. In the first five columns, each image is an averaged result across four scans for each subject. The image in the sixth column is the average of the results of all previous five columns.

The functional connectivity map estimated by p-correlation for the shuffled data in Fig. 1 (left panel) and for the phase-randomized data in Fig. 2 (b) are further compared with the correlation results in Figure S3 and S4.

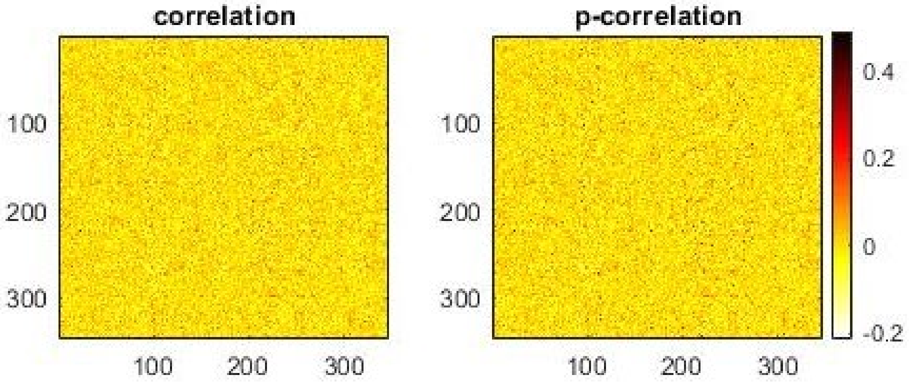


Figure S3 The functional connectivity map estimated by correlation (left) and p-correlation (right) on shuffled data. The correlation map has a mean of $0.0001 \pm 0.0402$. The p-correlation map has a mean of $0.0006 \pm 0.0414$. The two matrices are displayed in the same colormap

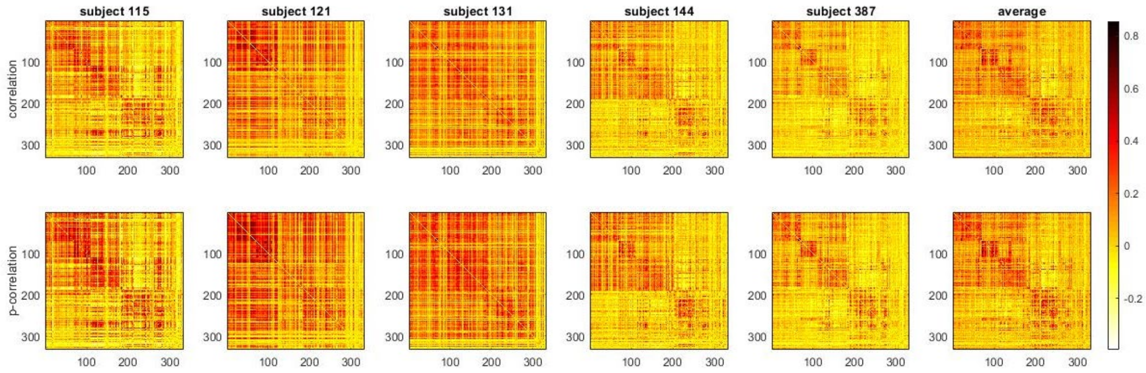


Figure S4 The connectivity map estimated by correlation (top) and p-correlations (bottom) on phase randomized data for five randomly selected subjects. In the first five columns, each image is an averaged result across four scans for each subject. The image in the sixth column is the average of the results of all previous five columns. The average correlation map across all scans and all subjects has a mean of $0.0737 \pm 0.1207$; the total average of p-correlation map has a mean of $0.0820 \pm 0.1253$.

### 2. Network determination and the network interaction matrices

Seven functional networks were determined by merging the 12 networks provided in Gordon, Laumann et al. (2016), into the 7 networks provided in Yeo, Krienen et al. (2011), through comparing their MNI coordinates. Specifically, the "Auditory", "Somatomotor Hand (SMhand)", "Somatomotor Mouth (SMmouth)" networks in Gordon, Laumann et al. (2016) become the new Somatomotor (SM) network. The "Ventral Attention", "Cingulo Opercular", "Cingulo Parietal" and "Salience" networks in Gordon, Laumann et al. (2016) are grouped into the new Ventral network. The new Default network includes the “Retrosplenial Temporal” and “Default” networks in Gordon, Laumann et al. (2016). The Frontoparietal, Visual, Dorsal and Limbic networks are the same as the ones in Gordon, Laumann et al. (2016). The seven networks were further verified by Informap graph analytical algorithm (Lancichinetti and Fortunato 2009), which returned roughly 7 disjoint groups among the 333 ROIs.

To evaluate the functional interactivity among networks, the functional connectivity map and the duration map were further partitioned into $S \times S$ subblocks, in which the diagonal subblocks includes the connectivity within functional networks and the off-diagonal subblocks includes the connectivity between different pair of networks. The average which includes the mean value and the standard deviation of each subblocks was computed, which results in four $S \times S$ matrices, namely the network interaction matrices. Figure S5 shows the network interaction matrices, which includes the mean value and the standard deviation of the connectivity strength and of the duration between every pair of networks. The numerical values in the four matrices of are tabulated in Table S1 and S2 of Supplementary Methods and Results.

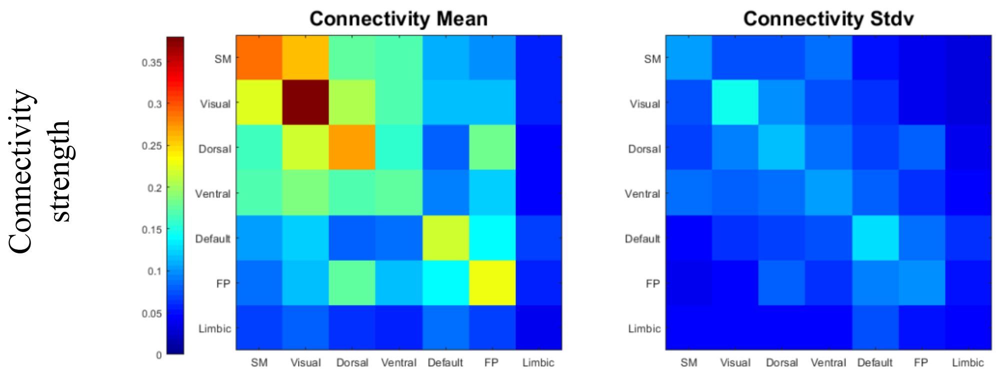

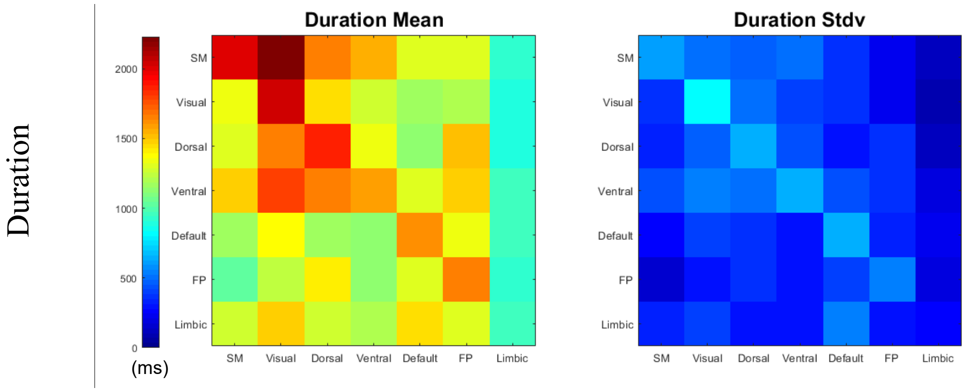


Figure S5 Network interaction matrices for the connectivity strengths and durations. The seven brain functional networks, SM, Visual, Dorsal, Ventral, Default, FP and Limbic are listed in four matrices. Each diagonal block shows the mean (left) and the standard deviation (right) of the connectivity strengths (upper) and of the duration of the information transfer (bottom) within each of the seven networks. Each off-diagonal block shows the average interactions (including the outward (the lower off-diagonal block) and the inward (the upper off-diagonal block) information transfer) between each of the 7 networks and one other network. The mean and standard deviation share the same colormap as shown in the left.

| **Connectivity Strength** | SM | Visual | Dorsal | Ventral | Default | FP | Limbic |
|---|---|---|---|---|---|---|---|
| SM | .287±.106 | .256±.077 | .174±.075 | .168±.087 | .107±.050 | .096±.036 | .055±.034 |
| Visual | .225±.073 | .377±.147 | .207±.095 | .169±.074 | .117±.062 | .117±.039 | .057±.032 |
| Dorsal | .163±.071 | .216±.095 | .273±.113 | .157±.088 | .080±.062 | .180±.077 | .046±.036 |
| Ventral | .168±.085 | .189±.078 | .168±.088 | .176±.106 | .094±.081 | .125±.062 | .046±.043 |
| Default | .103±.046 | .124±.061 | .082±.069 | .088±.076 | .218±.129 | .138±.087 | .070±.060 |
| FP | .088±.036 | .118±.045 | .177±.078 | .113±.063 | .137±.094 | .229±.096 | .055±.048 |
| Limbic | .068±.042 | .080±.043 | .060±.042 | .055±.047 | .088±.073 | .069±.053 | .038±.042 |

Table S1 Network interaction matrix for connectivity strengths. Mean values and standard deviations for the connectivity strengths of network interactions are shown.

| **Duration (ms)** | SM | Visual | Dorsal | Ventral | Default | FP | Limbic |
|---|---|---|---|---|---|---|---|
| SM | 2000±593 | 2235±514 | 1642±475 | 1549±505 | 1317±356 | 1310±223 | 937±114 |
| Visual | 1338±370 | 2042±837 | 1452±514 | 1272±415 | 1181±373 | 1209±244 | 881±94 |

| | | | | | | | |
|---|---|---|---|---|---|---|---|
| Dorsal | 1297±332 | 1666±481 | 1858±661 | 1336±443 | 1137±294 | 1532±379 | 903±123 |
| Ventral | 1495±436 | 1781±532 | 1648±522 | 1588±637 | 1319±421 | 1483±359 | 951±190 |
| Default | 1152±258 | 1362±413 | 1181±291 | 1121±636 | 1622±636 | 1330±320 | 970±214 |
| FP | 1040±170 | 1228±291 | 1423±287 | 1139±404 | 1300±404 | 1649±538 | 933±180 |
| Limbic | 1286±318 | 1494±389 | 1284±301 | 1187±292 | 1449±552 | 1312±290 | 953±267 |

Table S2 Network interaction matrix for durations. Mean values and standard deviations for the duration of network interactions are shown.

### 3. Short duration of information transfer in human brain

Because the limbic network has low functional connectivity (e.g., p-value>0.005 for p-correlations) due to the low temporal SNR of the fMRI timeseries, most information transfers with short durations that are within, coming into or coming out of the limbic network are not included in Figs. 2. The smallest 4% nonzero values in the duration map (without considering the p-value for p-correlations) and the corresponding connectivity map are shown in Figure and are jointly displayed on the brain cortex in Figure S6.

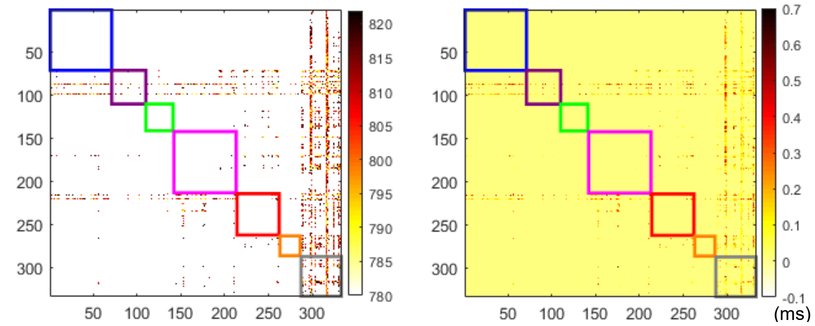


(a) Duration Map (b) Functional Map

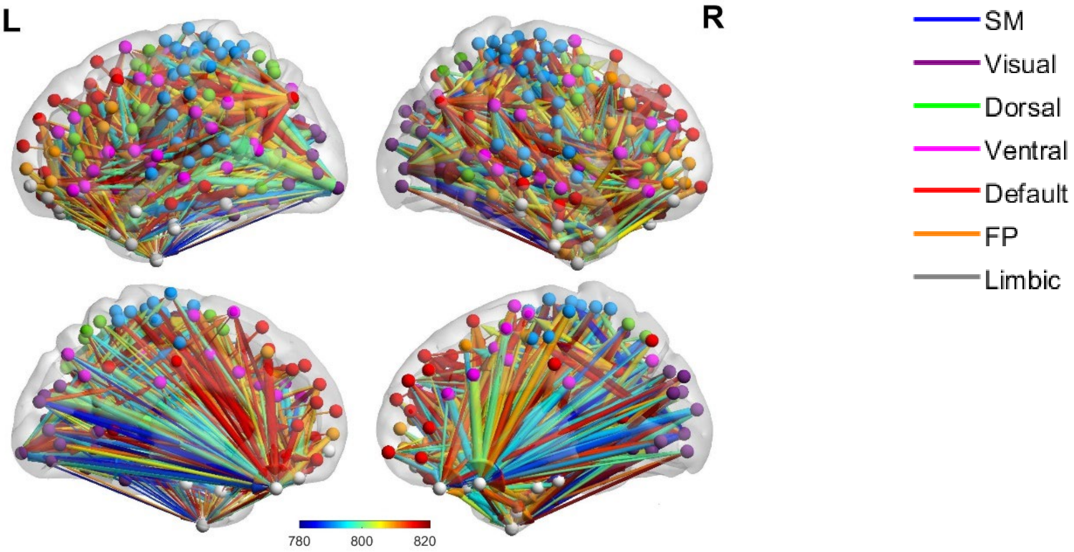


(c) The short information transfer on brain cortex

Figure S6 The short information transfer, characterized by the shortest 4% nonzero $N_{j|i}$ values. The duration map (a) and the connectivity map (b) of the short information transfer in resting brain. In both matrices, ROIs along the rows propagate information to the ROIs along the columns. Seven colored diagonal blocks in (a) and (b) depict the seven different networks as described in the legend (right). (c) The short information transfer on brain cortex. Stronger and longer connectivity has a thicker and warmer colored arrow. The node color corresponds the network as described in the legend (right).

### 4. Asymmetries between the inward and outward information transfer

| Direction / Network | (a) Within | (b) Inward | (c) Outward |
|---|---|---|---|
| Limbic | 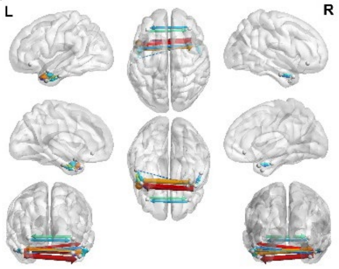  | 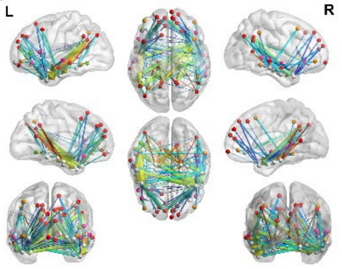  | 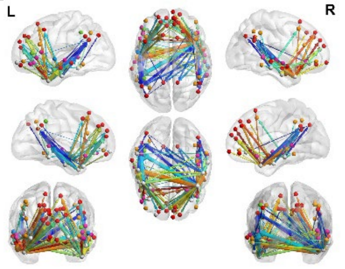  |
| SM | 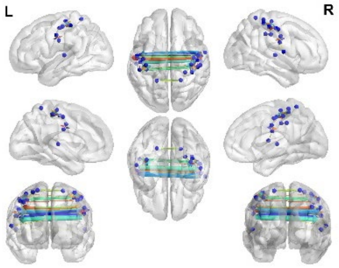  | 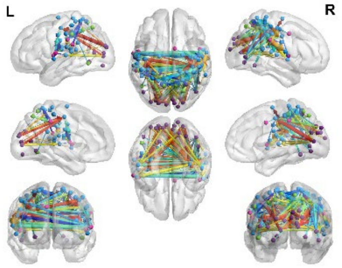  | 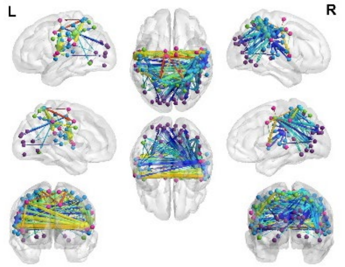  |

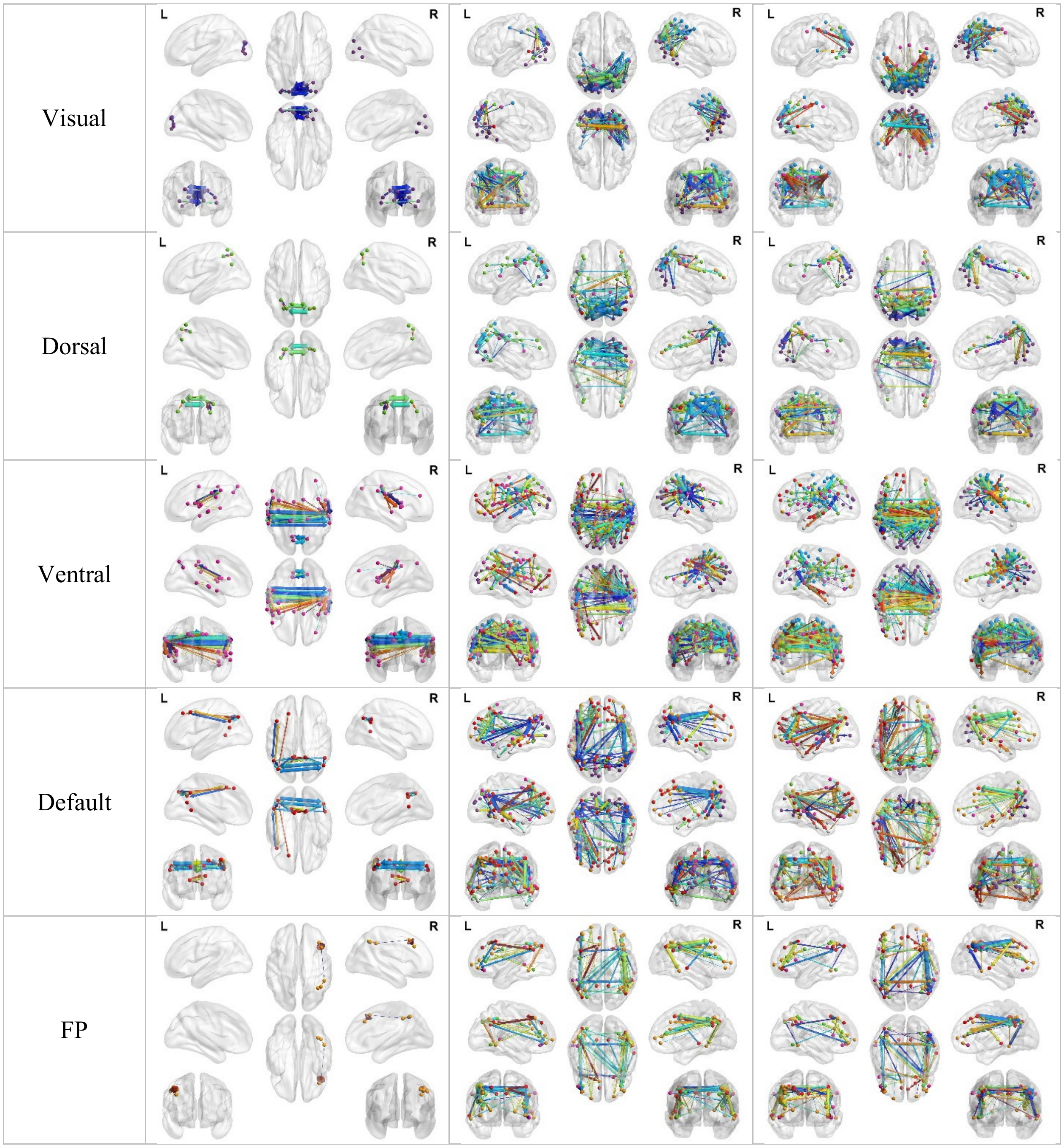


Figure S7 Spatial and Temporal interactions in resting state brain networks. The top 1% strongest connections within (a), coming into (b), and going out of (c) each of the seven functional brain networks are visualized by a modified version of BrainNet Viewer toolbox (Xia, Wang et al. 2013). Stronger and longer connectivity has a thicker and warmer colored arrow. ROIs in each of the six networks share the same network color as shown in the legend of Fig. 3 (right).
